# Bax and Bak jointly control survival and dampen the early unfolded protein response in pancreatic β-cells under glucolipotoxic stress

**DOI:** 10.1101/855239

**Authors:** Sarah A. White, Lisa Zhang, Yu Hsuan Carol Yang, Dan S. Luciani

## Abstract

ER stress and apoptosis contribute to the loss of pancreatic β-cells under the pro-diabetic conditions of glucolipotoxicity. Although activation of the canonical pathway of intrinsic apoptosis is known to require Bax and Bak, their individual and combined involvement in glucolipotoxic β-cell death have not been demonstrated. It has also remained an open question if Bax and Bak in β-cells have non-apoptotic roles in mitochondrial function and ER stress signaling, as suggested in other cell types. Using mice with individual or combined β-cell deletion of Bax and Bak, we demonstrated that glucolipotoxic β-cell death *in vitro* happens in sequential stages; first via non-apoptotic mechanisms and later by apoptosis, which Bax and Bak were redundant in triggering. In contrast, they had non-redundant roles in mediating staurosporine-induced β-cell apoptosis. We further established that Bax and Bak do not affect normal glucose-stimulated β-cell Ca^2+^ responses, insulin secretion, or *in vivo* glucose tolerance. Finally, our experiments revealed that Bax and Bak together dampen the unfolded protein response in β-cells during the early stages of chemical- or glucolipotoxicity-induced ER stress. These findings identify novel roles of the canonical apoptosis machinery in modulating stress signals that are important for the pathobiology of β-cells in diabetes.

## INTRODUCTION

In the pathogenesis of obesity-associated type 2 diabetes, chronic exposure to elevated blood glucose concentrations (glucotoxicity) and excess levels of circulating lipids (lipotoxicity) promote progressive failure and death of the insulin-secreting pancreatic β-cells (1-3). Moreover, the combined excess of glucose and lipids (glucolipotoxicity) may synergize to cause a faster and more severe progression of β-cell demise (4,5). Endoplasmic reticulum (ER) stress contributes to the loss of functional β-cells (1,6-8). ER stress activates the unfolded protein response (UPR), an adaptive measure initially aimed at restoring homeostasis through down-regulation of general protein translation and up-regulation of select genes encoding molecular chaperones and the machinery for ER associated degradation of proteins (ERAD). If the UPR fails to mitigate ER stress, a transition occurs whereby the UPR instead promotes apoptosis (9). While the involvement of ER stress in glucose- and lipid-induced β-cell pathobiology is widely accepted, many details regarding the regulation of the β-cell UPR, and its mechanistic links to β-cell death, remain unclear.

Chronic cellular stress triggers cell death by intrinsic apoptosis, which is regulated by proteins in the Bcl-2 family (10). Mitochondrial outer membrane permeabilization, a critical step in this process, is carried out by the pro-apoptotic family members Bax and Bak (11,12). Combined deletion of Bax and Bak thus provides a powerful means of dissecting the involvement of the canonical intrinsic apoptosis pathway. Unrelieved ER stress has also been demonstrated to trigger Bax and Bak-dependent death of several cell types (11,13-15). However, most studies of Bax and Bak in primary pancreatic β-cells under ER stress, including in human type 2 diabetes, have been correlative reports of increased expression or mitochondrial translocation of Bax (8,16,17). The few loss-of-function studies that have specifically addressed their requirement for β-cell apoptosis suggest that loss of either protein alone provides partial protection from cytokine-induced death (18), and that Bax^-/-^ β-cells show a modest protection from glucolipotoxicity (19). This could reflect significant overlap in signaling β-cell apoptosis, not unlike their redundant roles in cell death during development (20). However, apoptosis induced by chronic glucotoxicity or Pdx1 deficiency appears to preferentially engage Bax (21,22). This indicates that the relative involvement of the two proteins may be context- and stress-specific, but to date no study has included a full comparison of the individual and combined contributions of both Bax and Bak to stress-induced death in primary β-cells.

A growing body of work is also uncovering physiological functions of various Bcl family apoptosis proteins (23-32). We recently reported that anti-apoptotic Bcl-2 and Bcl-x_L_ modulate β-cell mitochondrial metabolism, glucose responsiveness, and ROS signaling (30,31). Others have shown that glucokinase activity is controlled by pro-apoptotic Bad (26,27). Studies in non-β-cells have implicated both Bax and Bak in ER physiology and UPR signaling (13,28), and Bax in the regulation of mitochondrial bioenergetics (29). It therefore remains an important question if Bax and Bak have non-apoptotic functions in pancreatic β-cells.

In this study we established lines of inducible knockout mice to determine the effects of single and combined Bax-Bak deficiency on β-cell function, ER stress signaling, and survival. We provide the first direct molecular assessment of the requirement for the intrinsic, mitochondrial, apoptosis pathway in glucolipotoxic β-cell death. Moreover, our data reveal that Bax and Bak do not play significant roles in β-cell function, but dampen the UPR in pancreatic islets during the early stages of ER stress.

## EXPERIMENTAL PROCEDURES

### Mice and in vivo studies

To establish the 4 genotypes of mice used for these studies we first bred Bax^flox/flox^:Bak^-/-^ mice (Stock number 006329, The Jackson Laboratory) with Pdx1-CreER mice (33) to create littermate mice with Bak single knockout (Bak SKO; Bax^flox/flox^:Bak^-/-^) and Bax-Bak double knockout (Bax-Bak DKO; Bax^flox/flox^:Bak^-/-^:Pdx1-CreER^+^) (34,35). We further mated Bax^flox/flox^:Bak^+/-^ progeny to re-introduce the wild-type Bak allele and obtain a parallel colony of Bax single knockout (Bax SKO; Bax^flox/flox^:Bak^+/+^:Pdx1-CreER^+^) and littermate wild-type mice (WT; Bax^flox/flox^:Bak^+/+^). Whenever possible, littermates were compared (Bak SKO vs Bax-Bak DKO, and Bax SKO vs WT). In all other instances age- and sex-matched mice from the two parallel lines were used. To induce Bax gene deletion, tamoxifen (3 mg/40 g body weight) was administered daily by intraperitoneal (ip) injection for 4 consecutive days. All mice, including controls, were similarly injected by tamoxifen. *In vivo* glucose tolerance was assessed by ip glucose tolerance tests after a 6 h fast. Glucose (2 g/kg body weight) was administered by ip injection and tail vein blood droplets were read using OneTouch Ultra Blue Test Strips (Lifescan) at the indicated time points. To minimize any putative long-term effects of Bax-Bak deletion, all *in vivo* experiments and islet isolations were done no later than 14 days following the final tamoxifen injection. The animal studies were approved by the University of British Columbia Animal Care Committee.

### Pancreatic islet isolation, dispersion, and culture

Mouse pancreatic islets were isolated from 12-16 week old mice by collagenase digestion followed by filtration-based purification, as previously described (30). Isolated islets were hand-picked and cultured overnight before further treatment. Palmitic acid was prepared in a 6:1 ratio with bovine serum albumin, as detailed previously (1). Unless otherwise indicated, islets were cultured in RPMI 1640 completed with 2% penicillin-streptomycin and 10% fetal bovine serum (Gibco, Thermo Fisher). For single-cell analyses, islets were dispersed into single cells prior to study, as described (36).

### Fluorescence Microscopy

Glucose-induced changes in cytosolic Ca^2+^ were compared in cultured intact islets using the ratiometric fluorescent Ca^2+^ indicator fura-2, as described (36). For measurements of mitochondrial membrane potential, dispersed islet cells were loaded for 30 minutes with 50 nM of the fluorescent indicator tetramethylrhodamine ethyl ester perchlorate (TMRE). Loading and imaging was done in phenol red-free RPMI 1640 and differences in mitochondrial membrane potential quantified by the absolute TMRE fluorescence intensity collected using a 585/60m emission filter following excitation using a 530/20x filter (Chroma Technology, Bellows Falls, VT, USA).

### Islet insulin secretion and content

For static assays of glucose-stimulated insulin secretion size-matched islets were first pre-incubated for 1 hour in Krebs-Ringer Buffer (KRB; 129 mM NaCl, 4.8 mM KCl, 1.2 mM MgSO_4_, 1.2 mM KH_2_PO_4_, 2.5 mM CaCl_2_, 5 mM NaHCO_3_, 10 mM HEPES, 0.5% bovine serum albumin) containing 3 mM glucose. The islets were then incubated in KRB with 3 mM glucose and then 20 mM glucose for 1 hour each, followed by 30 min incubation with KRB containing 3 mM glucose + 30 mM KCl. Supernatant was collected after each stimulation and assayed for secreted insulin. For quantification of insulin content, size-matched islets were washed twice with 1xPBS, collected in a 0.1 M HCl/70% ethanol solution and sonicated. Secreted insulin and insulin content were measured by radioimmunoassay (Rat insulin RIA Kit, Cedarlane, Burlington, ON, CA) or using the Mouse Ultrasensitive Insulin ELISA kit (ALPCO, Salem, NH, USA).

### Cell death assays

For quantification of cell death, dispersed islet cells were seeded into 96-well plates (Perkin Elmer ViewPlates) and allowed to adhere for 48 hours in complete media. As previously detailed (30), the cultures were stained with Propidium Iodide (0.5 μg/ml) and Hoechst 33342 (0.05 μg/ml) for 30 min before exposure to the indicated stress or control conditions. The 96-well plates were placed in an environmentally controlled (37°C, 5% CO_2_) ImageXpress Micro high content screening system (Molecular Devices) for the duration of imaging. Subsequently, cell death was determined as the number of PI positive cells relative to the number of Hoechst 33342 positive cells using the MetaXpress software (Molecular Devices, San Jose, CA, USA).

### Real-time PCR analysis of islet mRNA

Total islet mRNA was extracted using the RNEasy Mini Kit (Qiagen) and cDNA was synthesized by reverse transcription using 100 ng RNA and the qScript cDNA synthesis kit (Quanta Biosciences). Target gene expression was measured relative to mouse β-Actin housekeeping gene using PerfeCTa SYBR Green SuperMix plus ROX (Quanta Biosciences) and assayed using Applied Biosystems StepOnePlus and Applied Biosystems 7500Fast Real-Time qPCR machines. Primers were synthesized from IDT (Table 1).

**TABLE 1.**
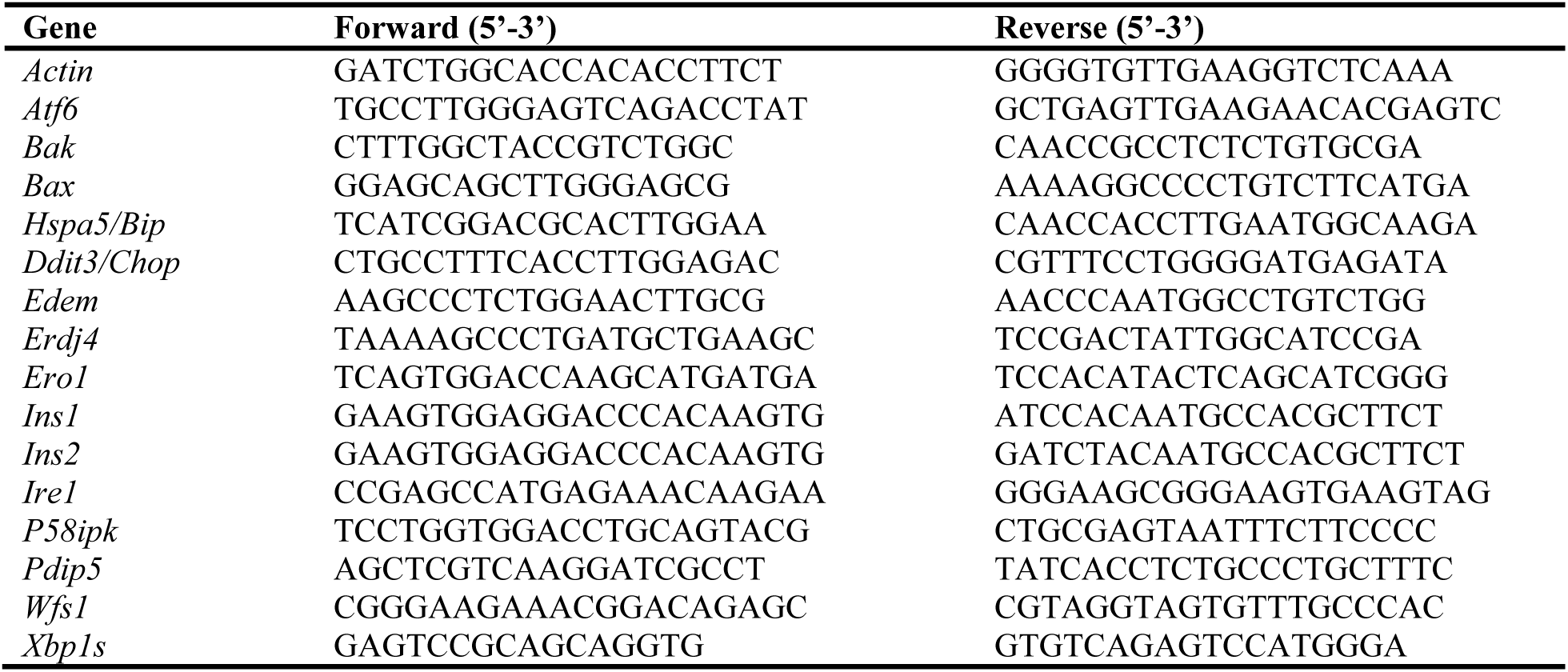
Quantitative PCR primer sequences.

### Reagents and compounds

InSolution™ Staurosporine was from Calbiochem/EMD Millipore. Dimethyl Sulfoxide (DMSO), Thapsigargin (Tg), Palmitic Acid (PA), Propidium Iodide (PI), Tetramethylrhodamine Ethyl Ester Perchlorate (TMRE), Collagenase Type XI from *clostridium histolyticum*, and Tamoxifen (TM) were from Sigma-Aldrich (St. Louis, MO, USA). Fura-2/AM and Hoechst 33342 were from Invitrogen/Life Technologies (Burlington, ON, CA).

### Statistical analysis

Data are presented as mean ± SEM. Data analysis was performed using GraphPad Prism 6.0 software. Statistical analysis was performed by student’s t-tests, 1-way ANOVA, or 2-way ANOVA with Bonferroni post-hoc comparisons where appropriate. Differences were considered significant if p < 0.05.

## RESULTS

### Loss of Bax and/or Bak does not affect normal islet function and glucose homeostasis

In mice, combined global knockout of Bax and Bak results in severe developmental abnormalities and perinatal death of most double knockout animals (20). Tissue-specific gene deletion is therefore required to study the effect of combined Bax and Bak loss. We previously established a line of Bak null mice in which the additional deletion of Bax can be selectively induced in β-cells of the adult animals by tamoxifen administration (30). To allow a comprehensive study of the individual and combined roles of Bax and Bak, we have now expanded this mouse colony to include the four genotypes: Wild-type (WT), Bax single-knockout (Bax SKO), Bak single knockout (Bak SKO), and Bax-Bak double knockout (Bax-Bak DKO) (cf. Experimental Procedures). Using real-time PCR we confirmed the complete absence of Bak mRNA, as well as the expected >80% knockdown of Bax in islets from all mice carrying the floxed Bax alleles following tamoxifen administration (Fig. 1). We previously demonstrated that the decreases in Bax and Bak mRNA are associated with similarly reduced islet protein (30).

**Figure 1.**
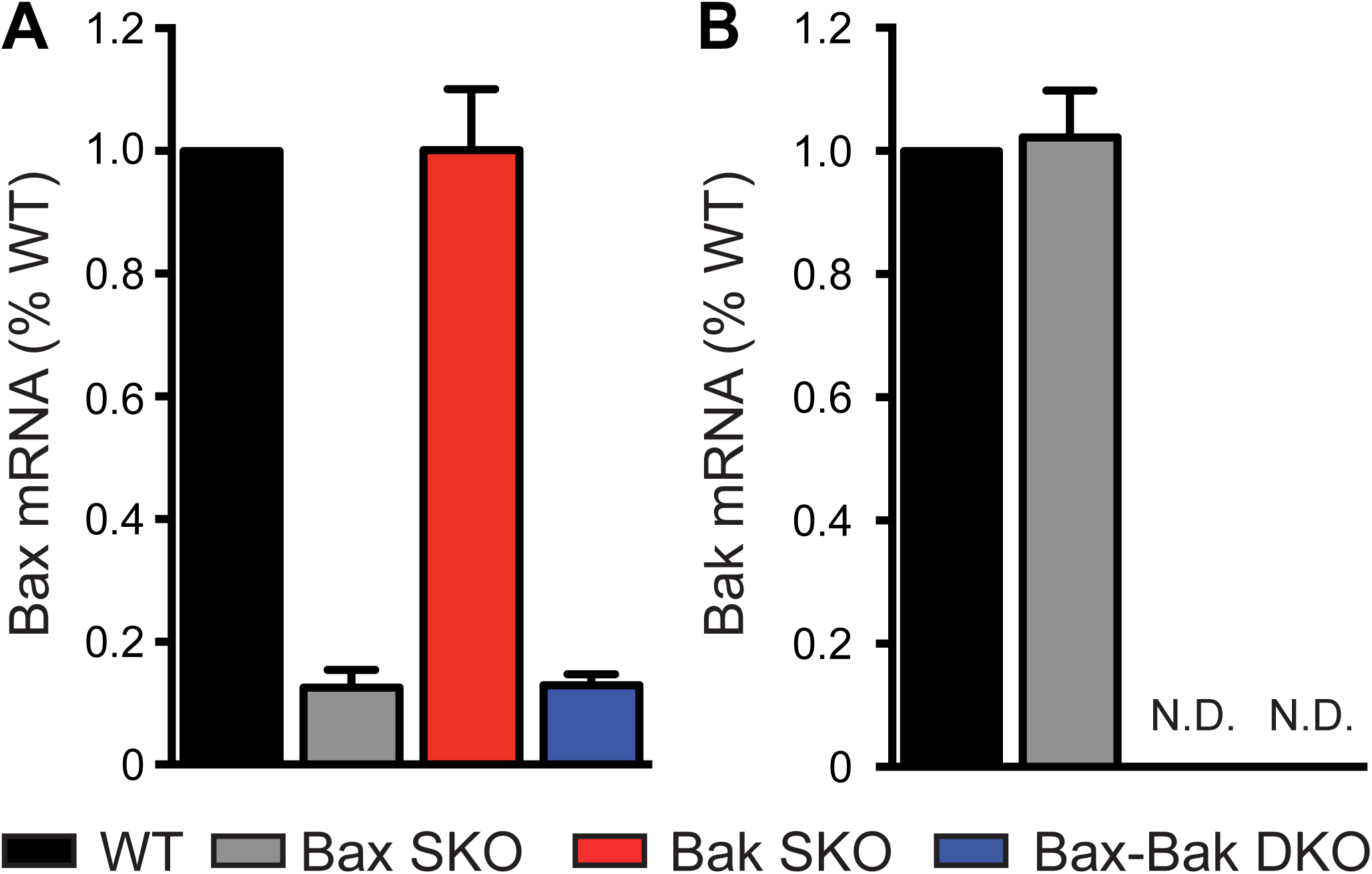
Combined and individual knockout of Bax and Bak in pancreatic islets. *A*, Relative Bax and *B*, relative Bak mRNA levels after tamoxifen administration and Cre recombination in islets from Bax SKO (Bax^flox/flox^:Bak^+/+^: Cre^+^), Bak SKO (Bax^flox/flox^:Bak^-/-^), and Bax-Bak DKO (Bax^flox/flox^:Bak^-/-^:Cre^+^) mice compared to WT (Bax^flox/flox^:Bak^+/+^) islets. (n= 4-6 animals of each genotype; islets isolated from 12-16 week old mice 1 week post-tamoxifen administration). Data represent mean ± SEM. N.D. = Not Detected.

Comparison of male and female mice of all four genotypes showed that single and double knockout mice all had normal body weights. Intraperitoenal glucose tolerance tests further demonstrated that there were no differences in the glucose tolerance of mice of all genotypes (Fig. 2). To more directly examine the impact of Bax-Bak deletion on β-cell function, we measured cytosolic Ca^2+^ responses and insulin secretion in isolated islets. No differences were observed in the cytosolic Ca^2+^ levels of WT and Bax-Bak DKO islets under basal conditions, or in the response to a stepwise glucose increase or direct depolarization with KCl (Fig. 3A,B). Double knockout of Bax and Bak did not affect islet insulin secretion under basal or acute glucose-stimulated conditions (Fig. 3C), and both islet insulin mRNA levels and insulin content did not differ between WT and Bax-Bak DKO islets in normal culture (Fig. 3D,E). Single deletion of Bax or Bak also did not affect islet function (data not shown). These results demonstrate that individual or combined loss of Bax and/or Bak in the adult β-cell has no detectable effects on the function of pancreatic islets under normal, non-stressed, conditions. Further, our comparison of the four different genotypes allows us to conclude that the life-long loss of Bak in all tissues does not affect *in vivo* glucose homeostasis.

**Figure 2.**
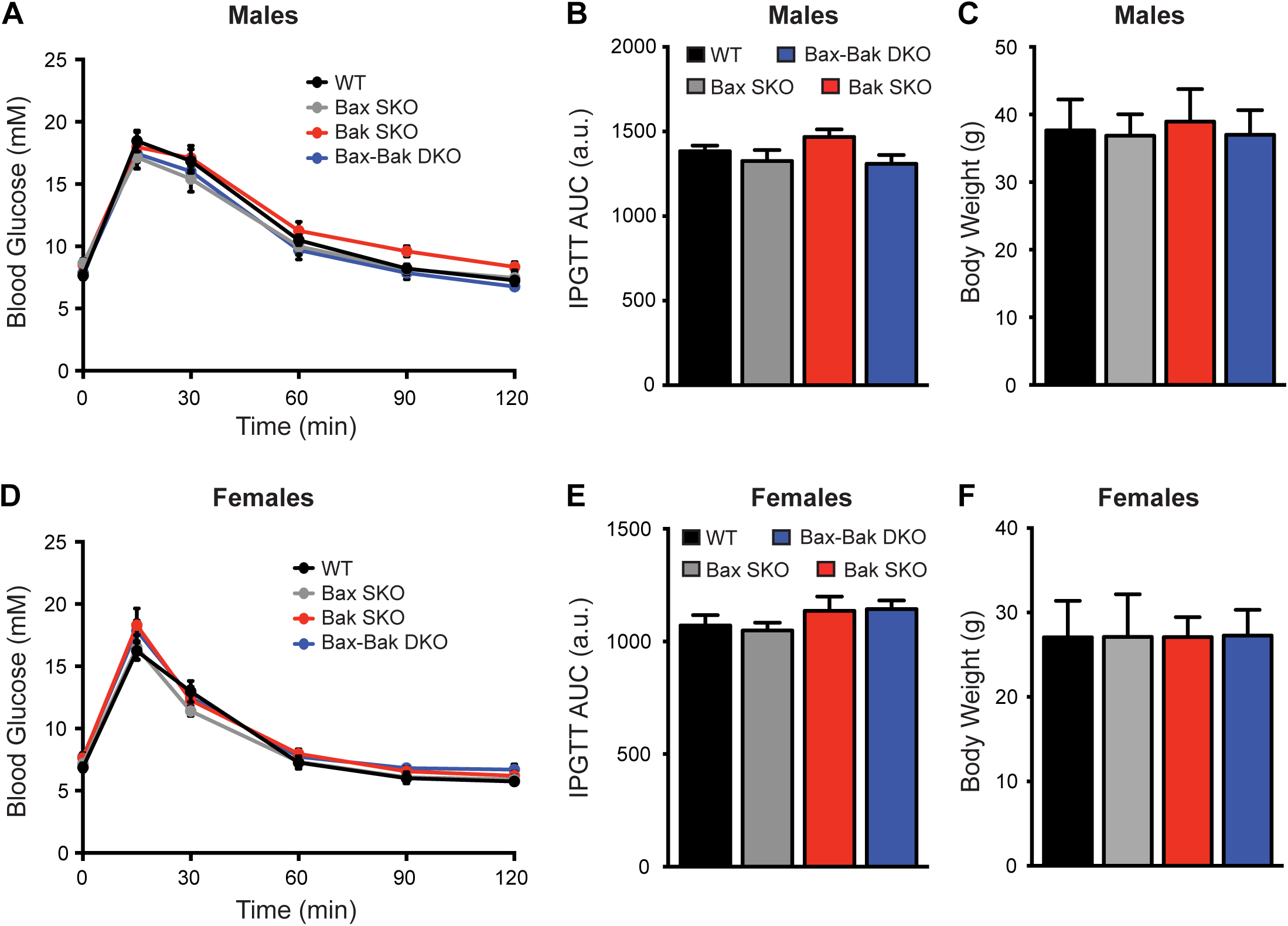
Loss of islet Bax and/or global Bak deletion does not affect body weight or *in vivo* glucose tolerance. Intraperitoneal glucose tolerance tests (IPGTT; 2 g/kg) and body weight analyses were performed in WT, Bax SKO, Bak SKO, and BaxBak DKO mice at 12-16 weeks of age. IPGTT blood glucose kinetics, area under the curve (AUC) of IPGTT profiles, and body weights, were not different between genotypes in male (*A-C*) and female (*D-F*) animals (n=15-30 animals of each genotype in each sex for body weight data; n=7-12 animals of each genotype in each sex for IPGTT data). Data represent mean ± SEM. a.u. = arbitrary units.

**Figure 3.**
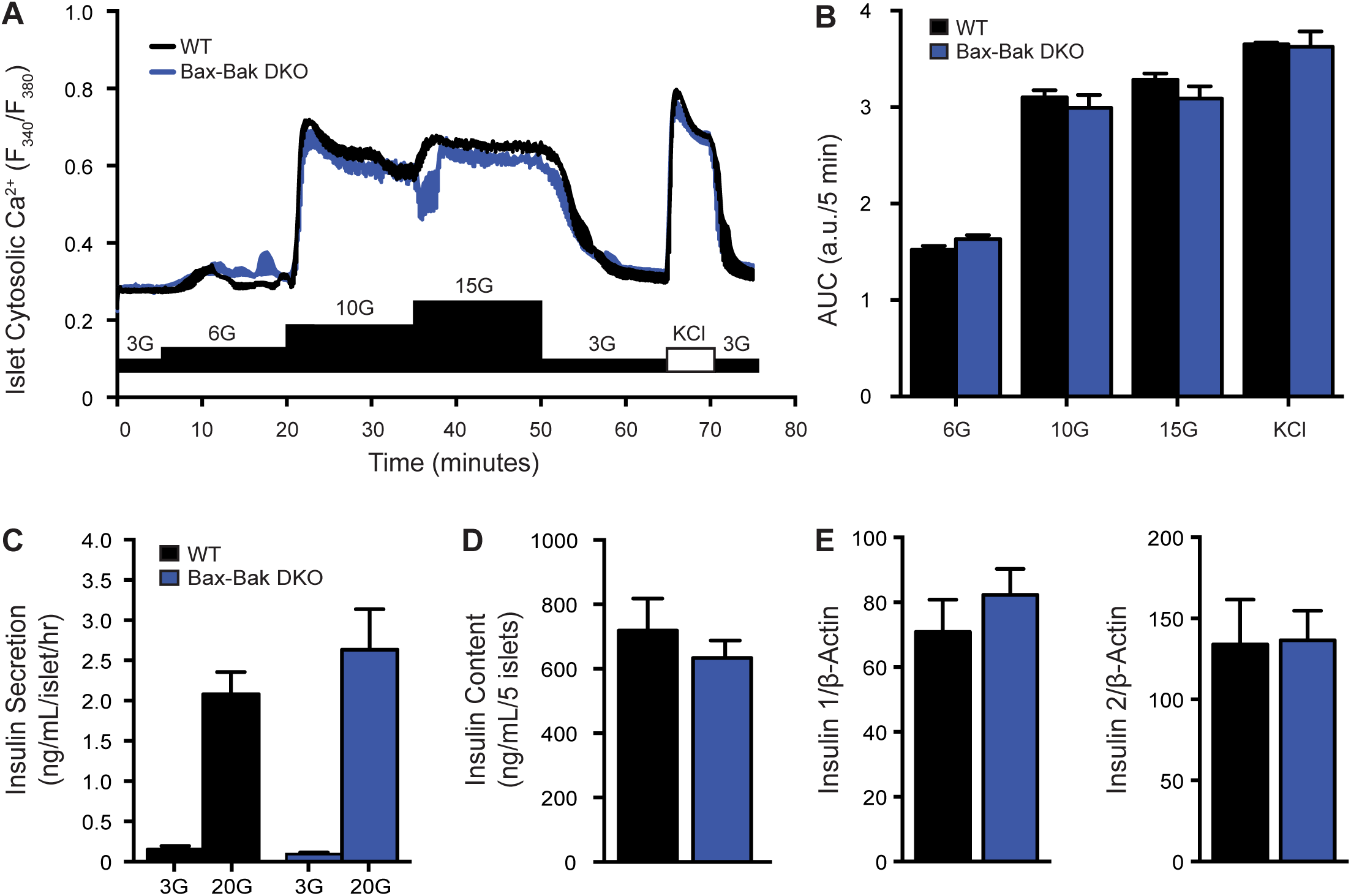
Loss of islet Bax and/or global Bak deletion does not affect islet physiology *in vitro*. *A*, Average cytosolic calcium (Ca^2+^) responses of whole islets from WT and Bax-Bak DKO mice stimulated sequentially with increasing glucose concentrations (G = mM glucose) and 3 mM glucose + 30 mM KCl (KCl). *B*, Area under the curve (AUC) analysis of the response to each stimulus in panel A. The AUC is expressed per 5 minute intervals for comparison (n=4 independent islet preparations). *C*, Glucose stimulated insulin secretion from static incubations of isolated WT and Bax-Bak DKO islets subjected to sequential stimulation for 1 hr each with 3 mM glucose (3G) and 20 mM glucose (20G) (n=7 independent experiments). *D*, Insulin content of size-matched islets from WT and Bax-Bak DKO mice (n=5 independent experiments). *E*, Insulin 1 and insulin 2 mRNA expression in WT and Bax-Bak DKO islets normalized to β-Actin housekeeping gene (n=4). Data represent mean ± SEM, a.u. = arbitrary units.

### Single and double Bax/Bak deletion reveals non-redundant roles in STS-induced β-cell apoptosis

To investigate the relative importance of Bax and Bak in executing β-cell apoptosis, we first performed a detailed kinetic analysis of staurosporine (STS)-induced death. STS is a pan-kinase inhibitor that has been demonstrated to activate Bcl-2-sensitive, Bax-Bak-dependent, apoptosis (11,37). Analysis of STS-induced death in islet cells of all four genotypes showed that WT cell death was initiated after roughly 12 hours and reached a maximal plateau by 36 hours (Fig. 4A). Absence of Bax and/or Bak markedly prevented cell death, with their combined deletion providing the highest degree of protection (Fig. 4A,B). Overall, the onset of STS-induced death of Bax SKO islet cells was delayed by several hours and then progressed at a significantly reduced rate. No death of Bak SKO and Bax-Bak DKO cells was detected until after more than 24 hours of STS-induced stress. That Bax SKO islet cells were less protected than those from Bak null mice likely reflects death of non-β islet cells that do not express Cre and therefore still have Bax. Consistent with loss of mitochondrial integrity in STS-induced apoptosis, WT islet cells showed a significant reduction in mitochondrial membrane potential after 24 hours. In contrast, the mitochondrial polarization of Bax-Bak DKO β-cells did not change, further supporting that mitochondrial outer membrane permeabilization was blocked at this time-point (Fig. 4C). Notably, islet cell death reached similar levels in all genotypes after approximately 48 hours of exposure to STS (Fig. 4A), indicating the activation of late-stage β-cell death that proceeds independently of the canonical machinery for intrinsic apoptosis.

**Figure 4.**
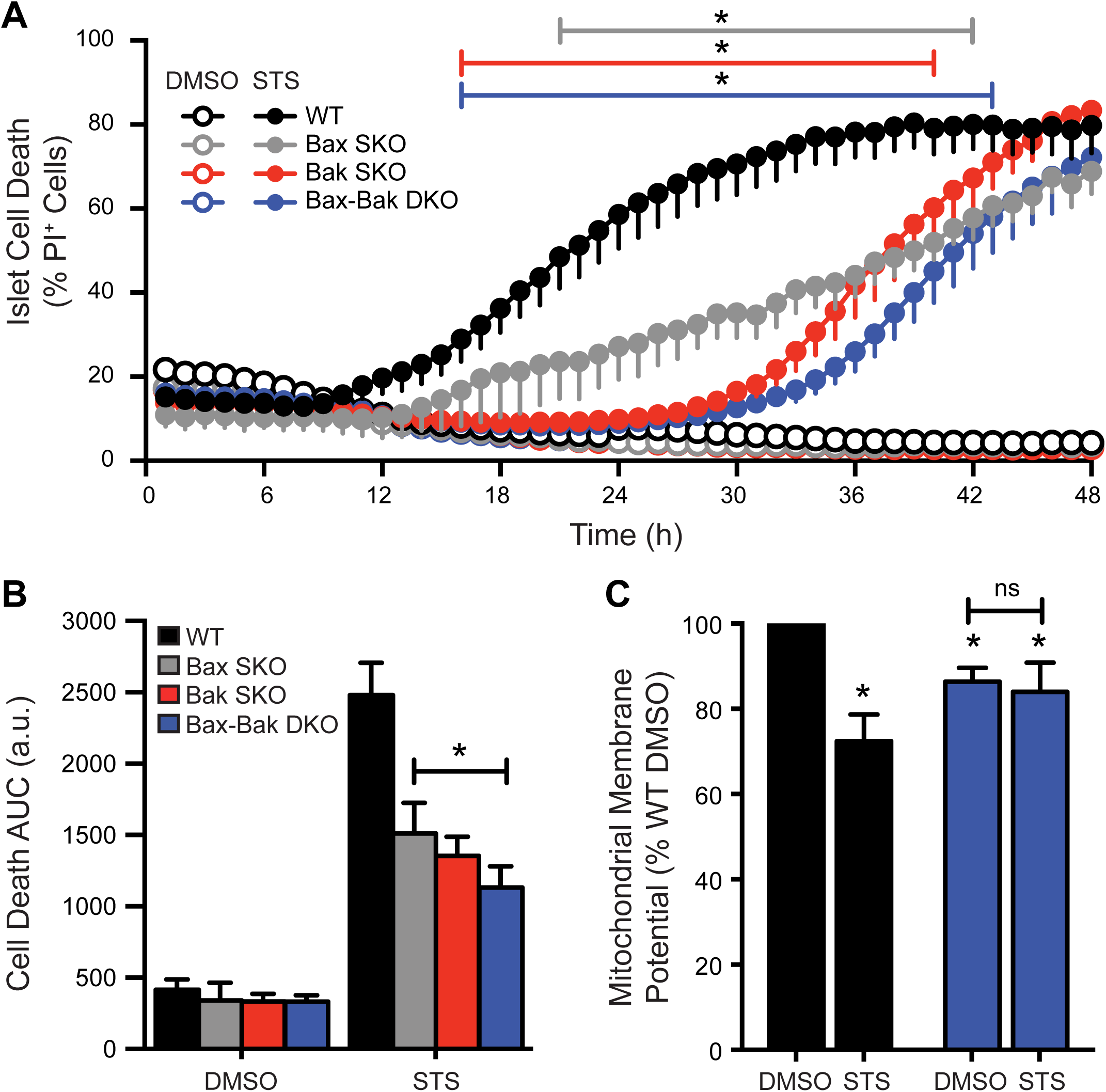
Relative contribution of Bax and Bak to staurosporine-induced mitochondrial apoptosis. *A*, Time-course cell death analysis of dispersed islet cells from WT (black), Bax SKO (gray), Bak SKO (red), and Bax-Bak DKO (blue) mice. Cells were treated with 1 μM staurosporine (STS; filled circles) or DMSO control (open circles) for 48 hours (h). The percentage of dead cells was measured as the number of propidium iodide positive (PI^+^) cells relative to total number of cells in each experiment. Coloured lines above the graphs indicate the time ranges where \**p* < 0.05 for the three knockout genotypes compared to WT. *B*, Area under the curve (AUC) analysis of the cell death profiles in panel A, \**p* < 0.05 compared to WT (n=4-11 independent islet cell preparations for the various genotypes). *C*, Mitochondrial membrane potential (Mean TMRE fluorescence intensity) of WT and Bax-Bak DKO islet cells treated with DMSO control or 1 μM STS for 24 hours. Data are normalized to WT DMSO control. \**p* < 0.05 compared to WT DMSO (n= 4 independent islet cell preparations). Data represent mean ± SEM, a.u. = arbitrary units.

Together, these findings demonstrate that the combined functions of Bax and Bak are required for STS-induced apoptosis in pancreatic β-cells, revealing that they cannot compensate for each other in this context. Further, our results show that chronic *in vitro* β-cell stress can eventually activate alternate, non-apoptotic, death mechanisms.

### Combined deletion of Bax and Bak is required for protection, and reveals multiple β-cell death modes, under glucolipotoxic stress

We next examined the requirement for Bax and Bak in mediating β-cell death during the diabetes-relevant conditions of glucose- and lipid-induced stress. Dispersed β-cells were cultured for up to 58 hours in the presence of varying levels of glucose with or without addition of the free fatty acid palmitate. Culture in low glucose (5 mM), high glucose (25 mM), or under strong lipotoxic conditions alone (5 mM glucose with 1.5 mM palmitate in 6:1 ratio with BSA), did not result in detectable death of islet cells from WT, Bak SKO, Bax SKO, or Bax-Bak DKO mice (Fig. 5 and data not shown). In contrast, glucolipotoxic stress caused progressive death of islet cells of all four genotypes (Fig. 5). Only the combined knockout of Bax and Bak provided significant protection against the glucolipotoxic insult, and this protection did not manifest until later stages of the experiment (Fig. 5A,B). Close inspection of the kinetic profile revealed a notable inflection point specifically in the death curve of Bax-Bak DKO cells at approximately 32 hours. Quantitation of the slope during 14 hour intervals before and after this time-point confirmed that the rate of death of WT and Bax-Bak DKO islet cells was similar at first, but then decreased significantly in the Bax-Bak DKO cells only (Fig. 5C). This suggests a time-dependent switch from predominantly non-apoptotic β-cell death toward Bax- and Bak-mediated apoptosis during chronic glucolipotoxic stress. Intriguingly, the comparison of all four genotypes further suggests that Bax and Bak are functionally redundant in signaling this late-stage β-cell apoptosis, which is in contrast to their combined requirement for STS-induced apoptosis.

**Figure 5.**
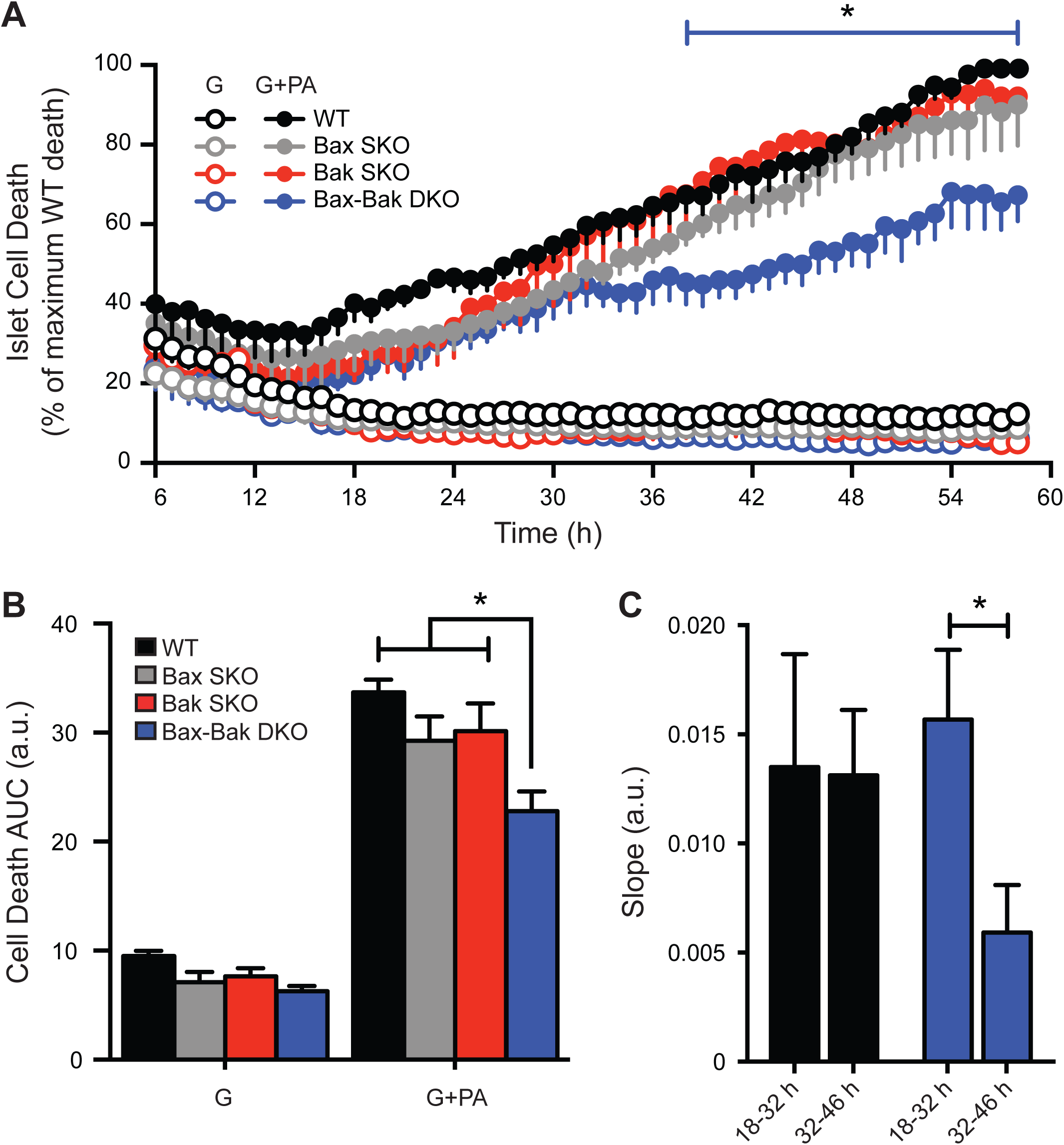
Combined knockout of Bax and Bak is required for significant protection against glucolipotoxicity. *A*, Time-course analysis of cell death (PI incorporation) in cultures of dispersed WT, Bax SKO, Bak SKO, and Bax-Bak DKO islet cells treated with 25 mM glucose (G; open circles) or 25 mM glucose + 1.5 mM palmitate (G+PA; closed circles) for 58 hours (h). Between experiments the degree of glucolipotoxic death after 58 hours showed some variability (15-25% between islet cell preparations), but was always lowest in the Bax-Bak DKO cells. To facilitate analysis, the islet cell death was expressed relative to that in the WT cultures at the end of each individual experiment (100%). The blue line above the graph indicates the time range where \**p* < 0.05 for Bax-Bak DKO cell death compared to WT, Bax SKO, and Bak SKO. *B*, Area under the curve (AUC) analysis of the cell death profiles in panel A \**p* < 0.05 for Bax-Bak DKO compared to WT, Bax SKO, and Bak SKO. *C*, Rates of WT and Bax-Bak DKO islet cell death were compared by calculating the slope of cell death profiles in panel A for two 14 hour intervals representing early (18-32 h) and late (32-46 h) stages. \**p* < 0.05 (n=4 independent islet cell preparations). Data represent mean ± SEM, a.u. = arbitrary units.

### Bax and Bak suppress the early unfolded protein response in β-cells under ER stress

Chronic hyperglycemia and elevated free fatty acids disrupt ER homeostasis and trigger ER stress, which contributes to β-cell dysfunction and death in diabetes pathogenesis. Motivated by evidence for involvement of Bcl-2 family proteins in the control of ER physiology (13,28), we next investigated whether Bax and Bak have roles in regulating β-cell UPR signaling under glucolipotoxic conditions. During ER stress, the expression of CHOP and XBP1s increases as a result of PERK and IRE1α activation, respectively. Under normal culture conditions WT and Bax-Bak DKO islets had similar expression levels of both CHOP and XBP1s (data not shown), indicating that Bax and Bak are not required for maintenance of basal β-cell ER homeostasis. Relative to culture in high glucose alone, the combination of high glucose and palmitate time-dependently increased CHOP and XBP1s mRNA in islets of all 4 genotypes (Fig. 6A,B and data not shown). Compared to WT, XBP1s transcripts were induced at significantly higher levels in Bax-Bak DKO islets after 24 hours, but this difference evened out by 48 hours (Fig. 6A). A trend toward higher CHOP mRNA was also seen in DKO islets after 48 hours, but this did not reach statistical significance (Fig. 6B). The individual deletion of Bax or Bak did not result in detectable differences compared to WT (data not shown). This suggests that Bax and Bak have redundant roles in dampening UPR signaling in β-cells under glucolipotoxic stress.

**Figure 6.**
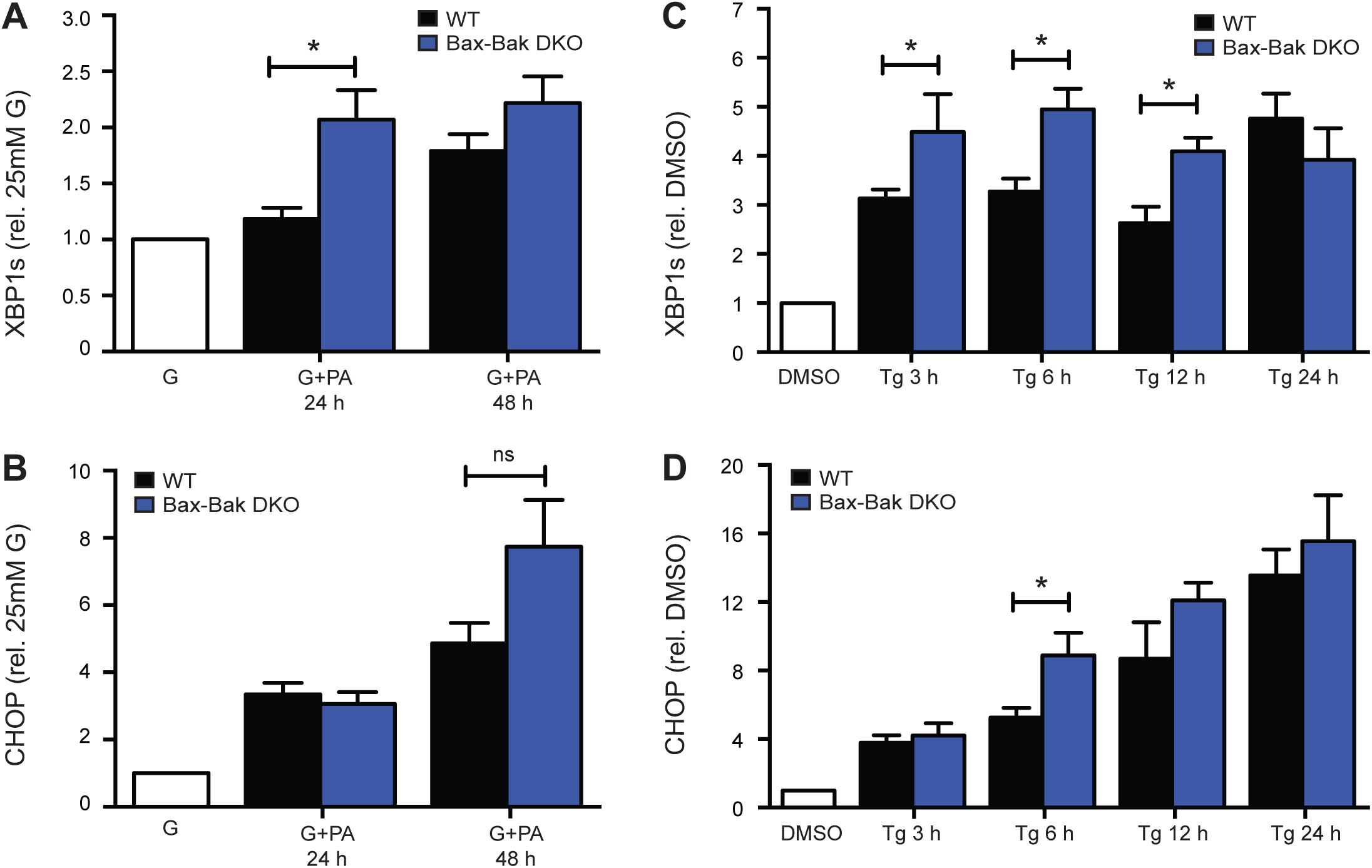
Double knockout of Bax and Bak augments early UPR signaling in islets under ER-stress. *A,B* Quantification of the time-dependent expression of XBP1s (A) and CHOP (B) mRNA in WT and Bax-Bak DKO islets treated with 25 mM glucose + 1.5 mM palmitate (G+PA) relative to 25 mM glucose along control (G; white bar) (n=3-7 at the different time-points). *C,D* Comparison of the time-dependent XBP1s and CHOP expression in WT and Bax-Bak DKO islets treated with 100 nM Thapsigargin (Tg) compared to DMSO vehicle control (white bar) (n=4-16 at the different time-points). Data represent mean ± SEM, \**p* < 0.05.

A glucolipotoxic insult activates a multifaceted β-cell response characterized not only by ER stress, but also by oxidative stress and general perturbations in other organelles (5). We therefore also examined the UPR following more specific ER stress, induced by thapsigargin, a chemical inhibitor of SERCA-mediated ER Ca^2+^ uptake (38). As expected, thapsigargin induced time-dependent expression of CHOP and XBP1s in the islets, with significant increases after just 3 hours (Fig. 6C,D). Similar to what was seen during glucolipotoxic stress, thapsigargin-mediated induction of XBP1s and CHOP mRNA levels were accelerated as a result of Bax-Bak double deletion, and the difference evened out following more chronic stress. Together, these data show that Bax and Bak dampen earlier stages of β-cell ER stress signaling involving both the IREα and PERK arms of the UPR.

To further assess the impact of combined Bax-Bak deletion on β-cell UPR signaling we examined the time-dependent expression of other UPR genes, including molecular chaperones and downstream target genes involved in the regulation of ER redox status and protein degradation. Thapsigargin-induced ER stress increased the expression of all genes examined (Fig. 7). In no instances did the basal expression of the UPR-related genes differ between islets from WT and Bax-Bak DKO mice, and under no conditions tested were UPR-related transcripts higher in the WT islets. However, in Bax-Bak DKO islets the mRNA levels of IRE1 itself were significantly increased after 6 hours treatment, and at 16 hours the mRNA levels of the chaperone BIP/HSPA5, the ER-associated glycoprotein WFS1 (Wolframin) (39), and the protein disulfide isomerase PDI-P5, were significantly higher compared to WT islets (Fig. 7).

These findings demonstrate that Bax and Bak together attenuate the UPR in pancreatic islets, revealing functions in β-cell stress signaling prior to activation of mitochondrial outer membrane permeabilization.

**Figure 7.**
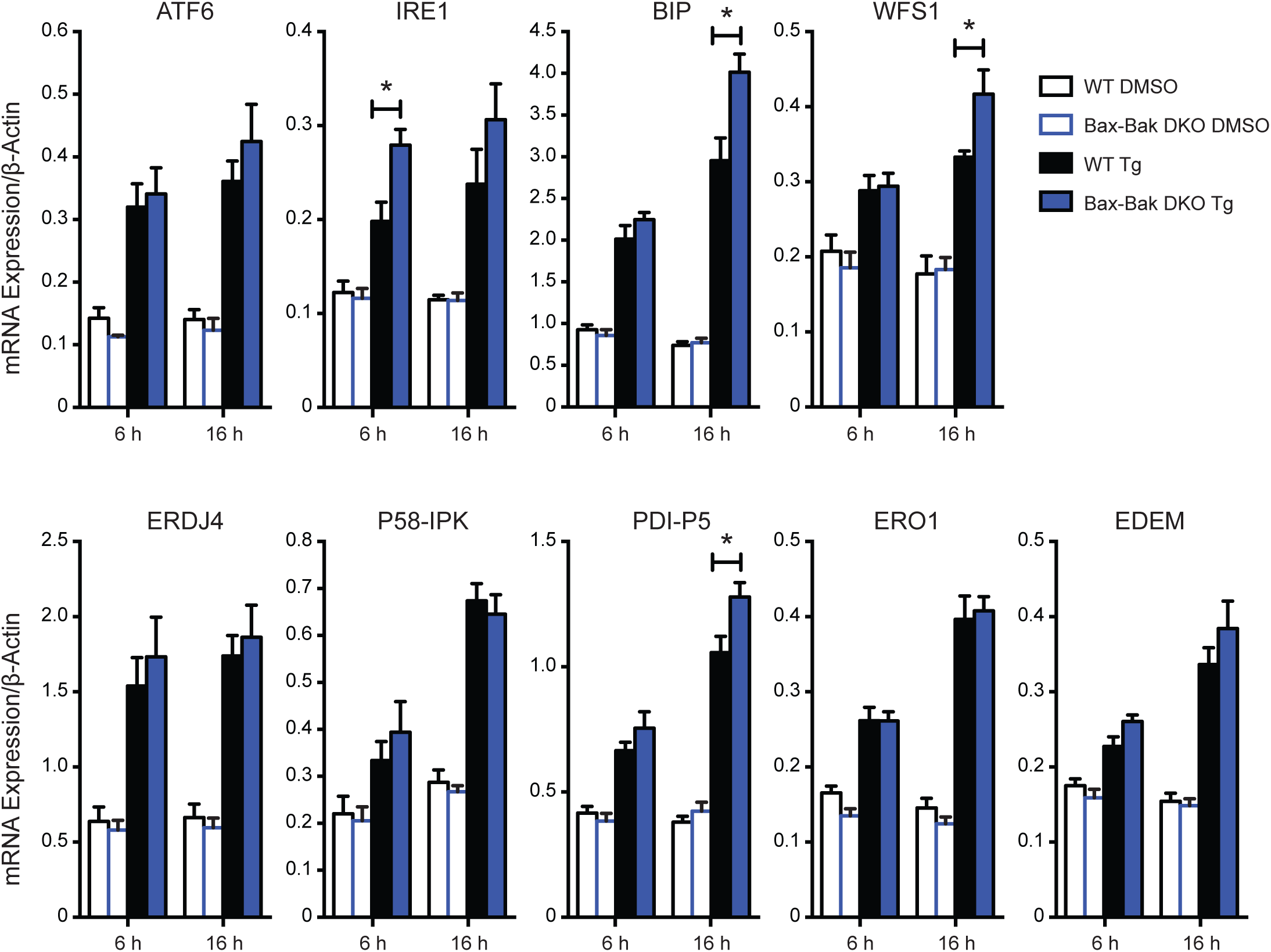
Downstream UPR gene expression in WT and Bax-Bak DKO islets under ER-stress. Gene expression analysis of UPR-inducers and downstream UPR target genes in WT and Bax-Bak DKO islets treated 6 or 16 hours with 100 nM Thapsigargin (Tg; coloured bars) or DMSO vehicle control (open bars). Gene expression was normalized to the housekeeping gene β-Actin. \**p* < 0.05 (n=6 independent islet preparations). Data represent mean ± SEM.

## DISCUSSION

In this study we used conditional and inducible gene deletion to dissect the roles of the apoptotic executioner proteins Bax and Bak in pancreatic β-cell function, as well as β-cell ER stress and death following combined exposure to high levels of glucose and fatty acids.

### Roles of Bax and Bak in the execution of β-cell death under diabetogenic stress

There is compelling evidence that apoptosis contributes to the loss of β-cell mass in human type 2 diabetes (3). In line with this, Bax mRNA and protein levels are significantly increased in islets from type 2 diabetic patients compared to non-diabetic controls (8,17). Islet levels of both Bax and Bak also increase during the progression of type 2 diabetes in the db/db mouse, but not in islets from obese ob/ob mice that remain normoglycemic (40). This suggests that Bax and Bak actively contribute to the diabetic islet phenotype. Accordingly, *in vitro* work has demonstrated roles for Bax, Bak, and other Bcl family members in β-cell death during lipotoxic and glucotoxic stress (16,17,21,41,42).

We did not detect β-cell death under culture with high glucose alone, or in response to palmitate in the presence of low glucose. This supports the idea that elevated glucose and lipids together induce a more detrimental ‘glucolipotoxic’ state (5,43). Others have demonstrated that glucotoxic β-cell death is Bak-independent, but involves Bax (21), likely via ‘upstream’ pro-apoptotic BH3-only family members Puma and Bim (12,17). This was observed using longer and more severe glucotoxic conditions than those we used here, and does therefore not disagree with our data. Lipotoxicity is known to activate Bax and β-cell apoptosis (1,16). The resilience of our primary mouse cells to pure lipotoxicity prevented us from clarifying the role of Bak in lipotoxic β-cell death.

Bax-Bak redundancy characterizes apoptosis during development (20) and also several known instances of pathogenic cell death *in vivo*, including of motor neurons in amyotrophic lateral sclerosis (44) and hepatocytes under severe ER stress (28). In this study, we found that primary β-cell death during the compound stress of glucolipotoxicity progressed normally with either Bax or Bak alone, suggesting a similar redundancy. In contrast, STS-induced β-cell apoptosis required the presence of both proteins. In conjunction with other studies that report preferential involvement of Bax in β-cell apoptosis under glucotoxicity (21) and Pdx1 deficiency (22), this illustrates that the relative requirement for Bax and Bak in β-cell death can depend on the specific form of stress.

Our Bax-Bak double knockout model allowed us to identify both apoptotic and non-apoptotic β-cell death during STS treatment and glucolipotoxicity. This agrees with the finding that stressed primary β-cells often die without all the morphological features of apoptosis (45). Moreover, death of Pdx-1-deficient β-cells happens by apoptosis, as well as necrosis involving the mitochondrial permeability transition pore (mPTP) (46-48). In myocardial infarctions deletion of Bax prevents both mPTP-driven necrosis and apoptosis (49), while loss of Bax in human leukemic CEM cells rather shifts ER stress-induced death from apoptotic to mPTP-dependent mechanisms (50). We observed late-stage death of STS-treated Bax-Bak DKO β-cells that might reflect an analogous apoptosis-to-necrosis switch. Inversely, our analyses indicated that non-apoptotic mechanisms dominate in the early stages of glucolipotoxic β-cell death. In future studies it will be important to further clarify the molecular nature, and putative crosstalk, of apoptotic and non-apoptotic β-cell loss under various stresses.

### Non-apoptotic roles of Bax and Bak in β-cell physiology and stress-signaling

Glucose-stimulated insulin release from β-cells depends on mitochondrial oxidative metabolism. A significant pool of cellular Bak is anchored in the outer mitochondrial membrane. In non-apoptotic cells this Bak binds to voltage-dependent anion channel 2, a protein involved in the mitochondrial transport of ions and metabolites (51). Furthermore, Bax and Bak have both been implicated in the control of normal mitochondrial fusion processes (52,53). This all hints that Bax and Bak might affect mitochondrial physiology. Indeed, Bax has been shown to promote mitochondrial bioenergetics in healthy HCT-116 cells and hepatocytes (29). Under normal culture conditions we observed a modest reduction in the mitochondrial membrane potential of Bax-Bak DKO islet cells, relative to WT. However, we established that this difference did not significantly alter glucose-stimulated Ca^2+^ signals and insulin secretion, which is further backed by normal glucose homeostasis in our knockout mice. In contrast, we and others have demonstrated that anti-apoptotic Bcl-2 and Bcl-x_L_ dampen β-cell glucose responses independently of Bax or Bak (30,54). Also, pro-apoptotic Bad modulates insulin secretion via interactions with glucokinase (27). It thus appears that Bcl-2 family proteins show different degrees of involvement in β-cell mitochondrial physiology.

Pools of Bax and Bak also localize to the ER (14), where they may affect cell survival via ER-derived Ca^2+^ signals (13,55). Previous work by Hetz *et al*. further found that loss of Bax and Bak impaired IRE1α-dependent UPR signaling in MEFs and hepatocytes (28). In pancreatic islets we observed the opposite; combined loss of β-cell Bax and Bak moderately amplified the early UPR under glucolipotoxic conditions and following chemical induction of ER stress. Since severe ER stress and chronic UPR signaling trigger β-cell apoptosis, it is possible Bax and Bak oppose their own activation. However, the UPR also serves an adaptive role (40,56). It is therefore also possible that Bax and Bak may impair the β-cell’s ability for ‘glucolipoadaptation’, thereby promoting both early dysfunction and late-stage death (43,57). Differentiating between these opposing outcomes will require very careful consideration of the time- and context-dependent transition from adaptive to apoptotic β-cell UPR signaling.

In summary, we used mouse models of inducible and conditional gene deletion to dissect the roles of pro-apoptotic Bax and Bak in β-cells. Our findings show that Bax and Bak do no play roles in normal β-cell function. Further, we provide new mechanistic insights into the mechanisms of glucolipotoxic β-cell loss and show for the first time that these pro-apoptotic Bcl-2 family proteins can modulate ER stress signaling in the endocrine pancreas.

## ACKNOWLEDGEMENTS

This work was supported by an operating grant from Diabetes Canada (OG-3-12-3787-DL) and D.S.L. received salary support from Diabetes Canada and the BC Children’s Hospital Research Institute (BCCHRI; Vancouver, BC). We thank Dr. James D. Johnson and Dr. Christopher A. Maxwell for generously providing access to their ImageXpress Micro high content imaging systems, and we would like to acknowledge Dr. Johnson for constructive input.

